# Checkpoint-dependent sensitivities to nucleoside analogues uncover specific patterns of genomic instability

**DOI:** 10.1101/2025.07.13.664472

**Authors:** Zainab Kagalwala, Mohammed Ayan Chhipa, Zohreh Kianfard, Essam Karam, Sirasie P Magalage, Sarah A Sabatinos

## Abstract

Nucleoside analogues are used as drugs and as labels in lab-based research. However, the effect of different nucleoside analogue mechanism(s) on cell sensitivity or mutagenesis is unclear. This is particularly important in cancer treatments where checkpoint proteins and DNA damage factors are often mutated. We tested 6 nucleoside analogues in the fission yeast, *Schizosaccharomyces pombe*. We found that the mutations in the DNA replication checkpoint cause unique sensitivity profiles towards chemotherapeutic nucleoside analogues (gemcitabine, 5-fluorouracil, cytarabine) and the non-clinical analogue bromodeoxyuridine. Antiretroviral compounds, zidovudine and lamivudine, did not alter cell growth. We compared half-maximal inhibitory concentration (IC50) doses between checkpoint deficient yeast strains, examining culture growth and DNA mis-segregation. Intriguingly, gemcitabine and bromodeoxyuridine doses above the IC50 promoted better growth. Above each compound’s IC50 dose we saw that cells were insensitive to nucleoside analogue re-exposure, particularly in DNA replication checkpoint mutants (*cds1*Δ*, rad3*Δ). Thus, pairing nucleoside analogue use with personal genomics may inform drug choice, dose, and schedule. Finally, these data indicate that resistance may be predictable, informing clinical strategy.

## 1. Introduction

Nucleoside analogues mimic DNA/RNA bases. The clinical uses of nucleoside analogues range from cancer chemotherapy to the treatment of Human Immunodeficiency Virus (HIV). Nucleoside analogue drugs work by a variety of mechanisms, often causing DNA replication inhibition and/or DNA damage *e.g*.[1–3]. Chemotherapeutics gemcitabine, cytarabine (AraC) and 5-fluorouracil (5FU) are antimetabolite prodrugs that are converted into active metabolites *in vivo,* and inhibit DNA replication in proliferating cells by incorporation, chain termination, and decreased dNTP levels [4]. Similarly, bromodeoxyuridine (BrdU) is incorporated into DNA during replication and can be detected using anti-BrdU antibodies [5]. BrdU is used to detect proliferation in lab studies [6, 7]. Antiretroviral nucleoside analogues, zidovudine (AZT) and lamivudine (3TC), inhibit viral replication through chain termination [8] and are an important component of HIV prophylaxis and treatment regimens. In all these cases, polymerase activity and proliferation are coupled to drug efficacy.

Because of the link between nucleoside analogues and DNA metabolism, exposed cells may arrest in the cell cycle. Cell cycle checkpoints regulate transition between phases of growth, DNA synthesis, and division in response to genome instability [9]. Checkpoint loss in cancer may make cells more sensitive to drug. Checkpoint proteins activate regulatory cascades that promote cell cycle arrest and repair [10–15]. Checkpoints prevent DNA under-replication, chromosomal changes, and mutations. Loss of checkpoints causes genome instability and induces cycles of DNA damage, chromosome mis-segregation, and aneuploidy *e.g*[16–20]. Mutational changes and genomic instability have a profound impact on the development of cancer in mammals *e.g.*[21–24], and development of heterogeneity within a population of cancer cells or tumour *e.g*.[21–23]. Cancer cells frequently mutate during drug treatment and adapt, causing a therapeutic obstacle through emergent drug resistance *e.g*.[25–27].

DNA replication arrest and DNA damage are two potent sources of genomic instability, carcinogenesis, and drug resistance [16, 20, 22, 26, 28]. DNA replication instability can be caused by a variety of environmental factors, including changes to nucleotide metabolism [29, 30] or polymerase inhibition [31–33]. Barriers including RNA-DNA R-loops and quadruplex DNA may block unwinding and replication fork movement [34–37], to cause DNA damage. DNA double strand breaks (DSBs) are considered a particularly lethal form of DNA damage that activate the G2-M DNA damage checkpoint [38–41]. DSBs are caused by ionizing radiation and some drugs (*e.g*. phleomycin) [38, 40–42]. DSBs are particularly dangerous if unrepaired by anaphase, causing genome loss or gain in daughter cells, chromosome fusions, telomere loss, and chromosome instability [43, 44].

Some nucleoside analogues are known to cause DNA damage and mutation. Yet, the link between checkpoint kinase function (genotype), initial drug sensitivity, and emergence of resistance is not well described. We used checkpoint-deficient mutants of fission yeast, *Schizosaccharomyces pombe*, to test nucleoside analogue sensitivity and deconvolve the relationships between dose and resistance. *S. pombe* is a rod-shaped, unicellular eukaryote that is phylogenetically distinct from the budding yeast *Saccharomyces cerevisiae* [45, 46]. Nucleotide metabolism, chromosome structures, and some DNA damage and repair mechanisms are more similar between *S. pombe* and humans than in budding yeast [6, 45, 47]. We tested single-gene loss of function mutations for nucleoside analogue effects, to compare with loss of checkpoint in human disease. Single mutations are stringently defined in our fission yeast model, which is not the case in mammalian cell lines that undergo mutations during transformation.

We developed a rapid proliferative assay to detect *S. pombe* culture proliferation and survival [48]. These results directly support nucleoside analogue studies in fission yeast, made by our group (Kianfard et al, *in prep*; and [6]) and others [7, 49–52]. We found that anti-cancer nucleoside analogues are most affected by loss of the DNA replication checkpoint (*cds1*Δ*, rad3*Δ mutants). Loss of the Rad3 apical checkpoint kinase (ATR orthologue; *rad3*Δ mutant) causes death at the half-maximal inhibitory concentration (IC50) dose. We propose that the IC50 dose is a “crisis point”, above which cell growth becomes less affected and potential for drug resistance increases. Anti-retroviral drugs do not affect cell proliferation; thus, AZT and 3TC do not induce a catastrophic checkpoint response. We hypothesise that increased sensitivity to nucleoside analogues in checkpoint mutant cells triggers DNA mis-segregation in replication checkpoint mutant cells (*cds1*Δ*, rad3*Δ). We conclude that nucleoside analogue chemotherapy treatments should be considered in the context of personal genomics, and that nucleoside analogues should not be paired with ATR checkpoint kinase inhibitors clinically.

## 2. Results and Discussion

*S. pombe* replication and damage checkpoints are initiated by the Rad3 kinase (**Figure 1**) [12, 53, 54]. Cells lacking *cds1*Δ or *rad3*Δ cannot activate the Intra-S and S-G2 checkpoints during replication stress; this causes DNA damage. Because the DNA replication checkpoint is important to processing nucleoside analogue effects, *cds1*Δ cells elongate, accumulate DBSs, and enter a lethal G2 arrest like in hydroxyurea [12, 13, 55]. Conversely, cells lacking *chk1*Δ can arrest in S-phase and stabilize DNA replication. However, *chk1*Δ cells cannot arrest in G2 if there is DNA damage [13, 14, 53]. Thus, *chk1*Δ cultures are a test for DSB-specific effects of nucleoside analogues. Because *rad3*Δ cannot arrest in either DNA replication instability or damage, we predicted that *rad3*Δ cells are most sensitive to nucleoside analogue effects. We expected that *rad3*Δ cells would present with DNA mis-segregation and “cell untimely torn” (*cut*) divisions, in which the septum forms over entangled DNA [56].

**Figure 1.**
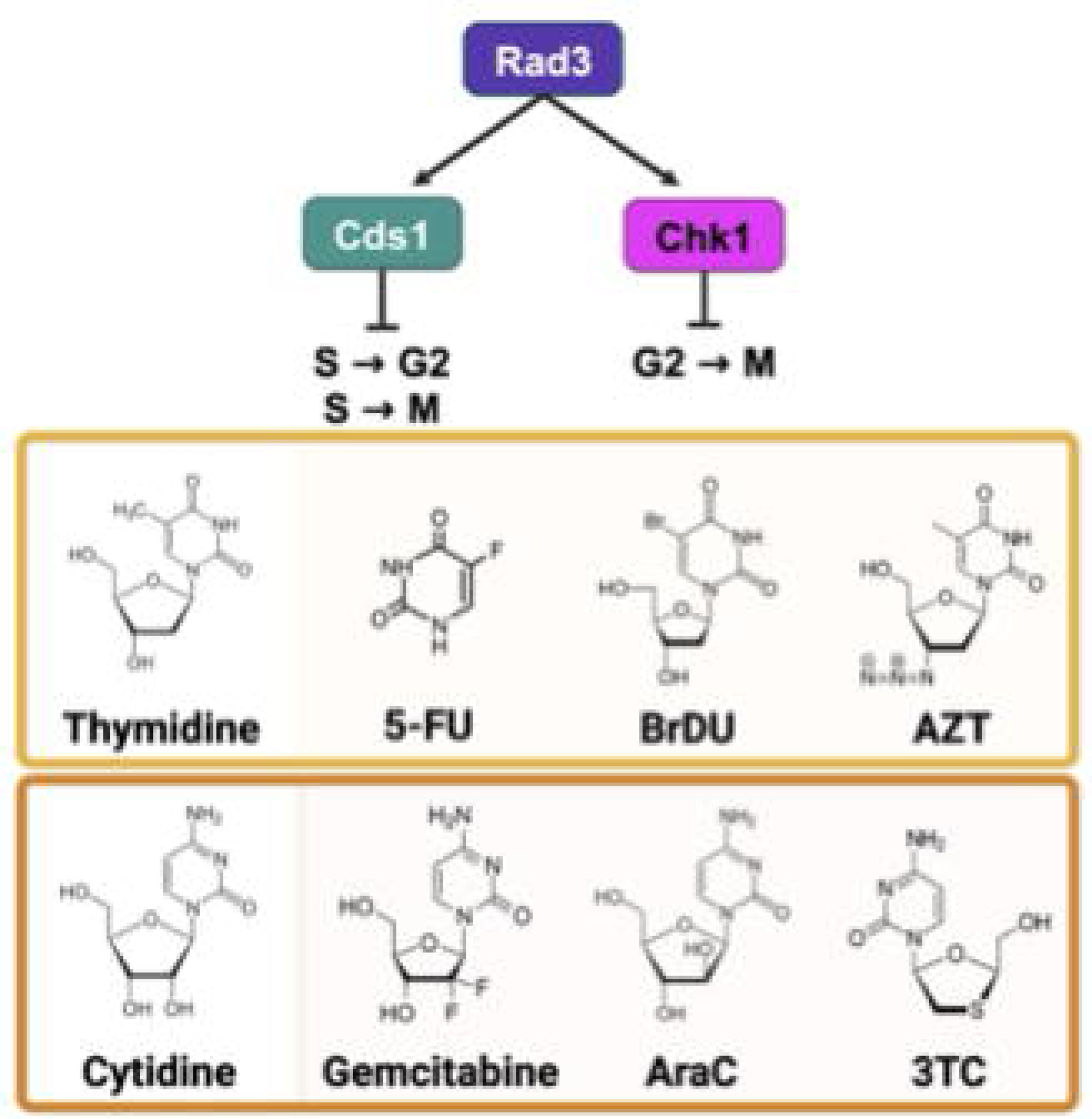
Checkpoint pathway and nucleoside analogues. A flow chart of the DNA replication (left) and DNA damage checkpoint pathways in *S. pombe.* Rad3 is an upstream kinase that is activated in response to both DNA replication instability and DNA damage to phosphorylate either Cds1 or Chk1. Below the checkpoint cartoon, chemical structures are shown. In the middle, thymidine is compared to its analogues 5’fluorouracil (5’FU), bromodeoxyuridine (BrdU), and zidovudine (AZT). The bottom row shows a chemical structure of cytidine and its analogues gemcitabine, cytarabine (AraC), and lamivudine (3TC).

We tested thymidine- and cytidine-like nucleoside analogues (**Figure 1B, 1C**) in *S. pombe* strains containing Herpes Simplex Virus (HSV1) thymidine kinase (*hsv-tk^+^*) and human equilibrative nucleoside transporter (hENT1+) transgenes (**Table 1**). Since *hsv-tk+* is a thymidine kinase, we hypothesized that thymidine-analogues would show the greatest effects. Because excess thymidine alters dNTP pools and causes S-phase arrest in mammalian cells [57] and fission yeast [6], we predicted that all thymidine analogues would arrest cell growth. BrdU causes mutation and cell death in *S. pombe* DNA replication checkpoint mutants [6]. We expected lower IC50 values for these *cds1*Δ and *rad3*Δ cells.

**Table 1.** Fission yeast strains.

| Strain | ID | Genotype | Source<sup>reference(s)</sup> |
| --- | --- | --- | --- |
| wild type (WT) | SY 17 | *h<sup>+</sup> leu1-32::hENT1-leu1+(pJAH29) his7-366::hsv-tk-his7+(pJAH31) ura4-D18 ade6-M210* | Susan Forsburg[38] |
| *cds1Δ* | SY 91 | *h<sup>+</sup> cds1Δ::ura4+ leu1-32::hENT1-leu1+(pJAH29) his7-366::hsv-tk-his7+(pJAH31) ura4-D18 ade6-M210* | Susan Forsburg[39] |
| *chk1Δ* | SY 92 | *h<sup>+</sup> chk1Δ::ura4+ leu1-32::hENT1-leu1+(pJAH29) his7-366::hsv-tk-his7+(pJAH31) ura4-D18 ade6-M210* | Susan Forsburg[39] |
| *rad3Δ* | SY 93 | *h<sup>+</sup> rad3Δ::ura4+ leu1-32::hENT1-leu1+(pJAH29) his7-366::hsv-tk-his7+(pJAH31) ura4-D18 ade6-M210* | Susan Forsburg[39] |

### 2.1. Thymidine analogues induce both DNA replication and DNA damage stress

We used our *S. pombe-*adapted dose-response methods to examine culture growth by OD600, and to calculate half-maximal inhibitory (IC50) doses for each analogue [48]. IC50 values allow comparisons between strain sensitivities and different drugs [48, 58]. IC50 values are used to suggest a drug dose that decreases proliferation [58]. Because *S. pombe* cells elongate during G2-checkpoint arrest, OD600 may increase simply because of bigger cells leading to higher turbidity. To determine if cells were truly dead or merely arrested, we spotted cultures onto drug-free medium after exposure.

We found that thymidine, 5 fluorouracil (5FU), and bromodeoxyuridine (BrdU) all decreased *cds1*Δ and *rad3*Δ mutant cell growth, while zidovudine (AZT) did not. Thymidine IC50 values were 16.8 µM for *cds1*Δ and 14.9µM for *rad3*Δ; however, thymidine did not decrease OD600 or generate an IC50 value in wild type or *chk1*Δ. A difference in growth was reflected in area under the curve (AUC), which was significantly lower in *rad3*Δ than wild type. The *rad3*Δ spot growth above 5 µM thymidine was slow with fewer cells per spot (Figure 2A iv). These data confirm that high levels of thymidine are toxic to *rad3*Δ cells. Cultures of *chk1*Δ cells had less spot growth at the highest thymidine doses. The fact that *cds1*Δ cells show similar IC50 curves in liquid growth indicate that thymidine causes DNA replication instability that relies on the DNA replication checkpoint for stability. However, because *rad3*Δ cells are more sensitive than *cds1*Δ, we infer that thymidine effects are not solely caused by replication checkpoint loss. Instead, *chk1*Δ growth reduction in spots after exposure to >100 µM thymidine indicates that high-dose thymidine causes genome instability possibly through DNA damage.

**Figure 2.**
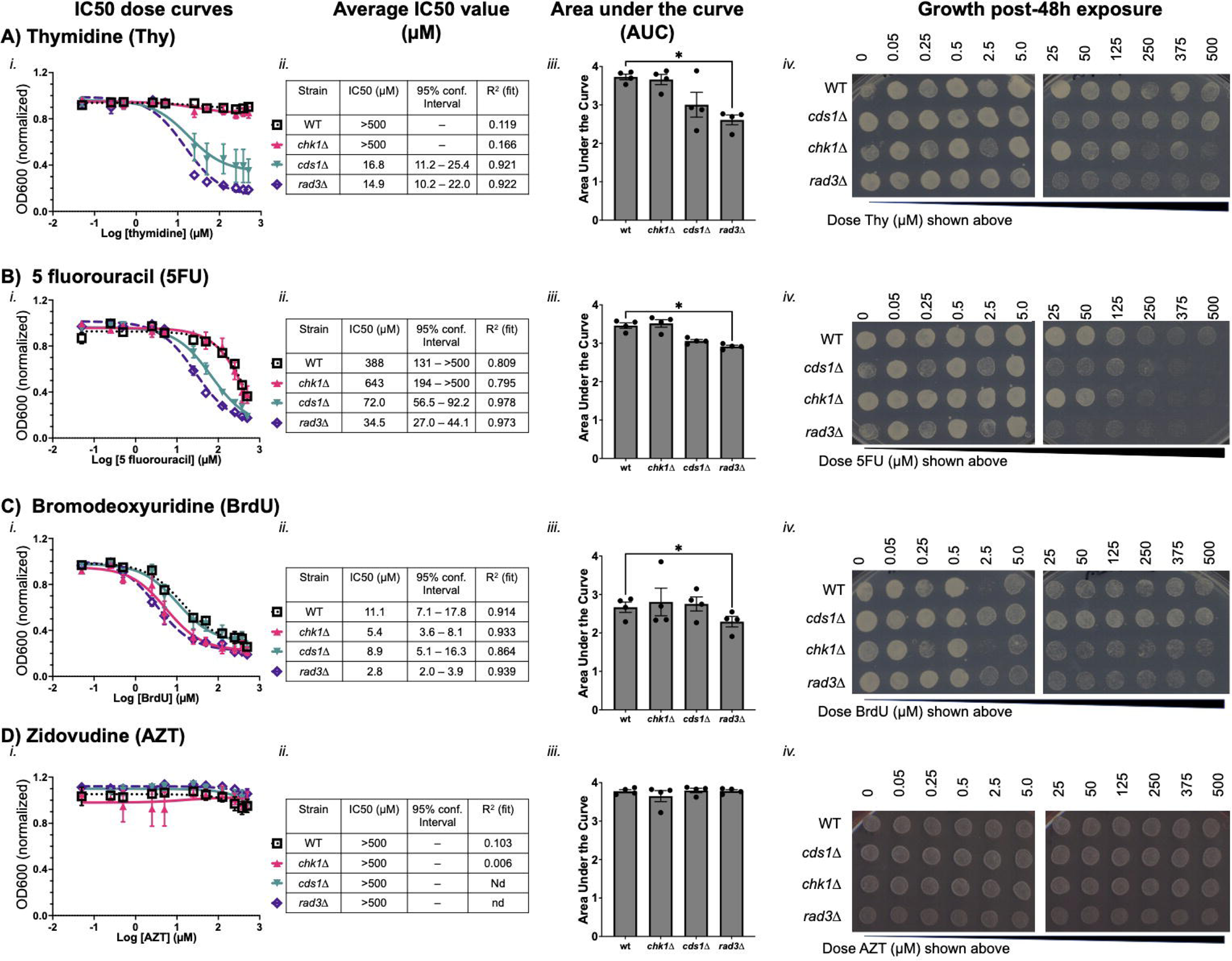
DNA replication checkpoint activity protects cells from thymidine analogue toxicity in fission yeast. A-D) Dose-response curves of S. pombe liquid culture growth. Optical density at 600 nm was used to measure proliferation. Shown are average points/curves with standard deviation from a minimum of 4 experimental replicates in A) thymidine (Thy), B) 5 fluorouracil (5FU), C) bromodeoxyuridine (BrdU), D) zidovudine (AZT). Average plots were used to model growth at doses of analogue and calculate half-maximal growth-inhibition (IC50) doses for each strain. The average IC50 value in µM is shown for each checkpoint mutant strain. Strains that were not sensitive have an IC50 value above 500 µM (>500, charts), which was the highest dose used. A 95% confidence interval and R^2^ value from 4 experimental replicates were calculated around the IC50 dose. In iii), The area under the curve (AUC) was calculated from a minimum of 4 experimental replicates and is plotted with standard deviation. A one-way ANOVA was used to compare AUC for each strain in each analogue (* p<0.05). iv) Cultures were pinned onto growth medium after 48h exposure to analogue. Plates were grown and photographed to detect whether an analogue caused growth arrest or cell death.

In contrast, 5-fluorouracil (5FU) and bromodeoxyuridine (BrdU) affected growth of all strains. 5FU was most potent in *rad3*Δ and *cds1*Δ (IC50s of 34.5 µM and 72.0 µM respectively) but 10-fold more 5FU is required to cause growth effects in wild type and *chk1*Δ (IC50s of 388µM and 643 µM). Spot growth after 48h of 5FU exposure is minimal in *rad3*Δ and *cds1*Δ above 5 µM (Figure 2B). Wild type and *chk1*Δ cells have reduced spot growth >50 µM. In all cases, the concentrations at which we observed reduced spot growth (Fig 2B iv) were lower than the calculated IC50s. These data indicate that 5FU effects require the DNA replication checkpoint to preserve viability. However, even replication-checkpoint competent cells are sensitive to high doses of 5FU, which impairs cell growth. Although *chk1*Δ spots appear no different than wild type after 5FU, the impact of DNA damage checkpoint loss should be investigated for differences in mutation kind and profile (i.e. [59]).

We found that BrdU also decreased OD600 and spot growth in all strains, and at lower doses than 5FU. BrdU is an important molecular tool to monitor DNA synthesis. Yet, the potential BrdU effects on DNA damage and proliferation are frequently overlooked in the interest of the desired experimental goal. We found that BrdU decreased turbidity in all strains, with IC50 values ranging from 2.8µM in *rad3*Δ to 11.1 µM in wild type (Figure 2C i, ii). AUC for *rad3*Δ was significantly different from wild type (p<0.05, Figure 2Ciii). While all spots showed diminished growth above 2.5 µM BrdU, *cds1*Δ spots were less affected by BrdU treatment (Figure 2C iv). Intriguingly, *chk1*Δ cells showed less growth by IC50 (5.4 µM) and in spots compared to *cds1*Δ. These data indicate that BrdU generates substantial DNA damage that may be more impactful than its effect on DNA replication stability. Because *cds1*Δ cells grew better than wild type after BrdU treatment, we conclude that loss of the DNA replication checkpoint in *cds1*Δ but presence of the DNA damage checkpoint may improve survival. In contrast, we infer that although OD600 turbidity is decreased in *rad3*Δ and *chk1*Δ, these cells acquire DNA damage and do not grow in the presence of BrdU. After spotting onto BrdU-free medium to grow post-exposure, the *rad3*Δ and *chk1*Δ cells re-enter growth slowly. We conclude that slower *rad3*Δ and *chk1*Δ growth is caused by DNA damage in BrdU which these G2/M checkpoint deficient strains cannot repair or arrest in the presence of.

AZT is an anti-retroviral polymerase inhibitor that did not inhibit growth in culture (Figure 2D i). All strains generated relatively flat dose-growth curves that cannot be used to calculate IC50 values. We reported all IC50 values as >500 µM, the highest dose that we tested. No significant difference in AUC values or spot growth was observed. Cells after 48h of AZT did not die but grew into full spots, even at 500 µM. We conclude that AZT is a non-toxic thymidine nucleoside analogue that does not impact growth during or after treatment, nor is this dependent on the DNA replication or damage checkpoints. This contrasts with 5FU, BrdU, and thymidine, all which arrest cell growth and cause long-term effects that decrease growth after exposure.

### 2.2. Cytidine anti-cancer analogues generate replication instability and not DNA damage arrest

Cytidine-related nucleoside analogue drugs include gemcitabine (Gem), cytarabine (AraC), and the anti-retroviral lamivudine (3TC). We used the same strains containing HSV1-TK. HSV1-TK is a promiscuous kinase that can phosphorylate a broad array of nucleosides including cytidine [38, 40]. Because altered CTP and dCTP levels result in mutagenesis, we hypothesized that HSV1-TK may cause genome instability effects from cytidine nucleoside analogue drugs. If so, cytidine-dependent DNA replication instability and/or DNA damage would be detected in specific checkpoint mutants. We predicted that replication checkpoint kinase mutant *cds1*Δ, or apical kinase *rad3*Δ, would show the least growth in cytidine analogues in our transgenic strains expressing hENT1^+^ and HSV1-TK.

We found that gemcitabine and AraC both decrease growth of replication checkpoint mutant cells. In this fission yeast-IC50 test of comparative dose-response effects, gemcitabine caused growth arrest in *cds1*Δ and *rad3*Δ cells (Figure 3A, IC50 values of 2.7 µM and 0.9 µM respectively). The area under the curve for *cds1*Δ and *rad3*Δ was also significantly lower than wild type cells in gemcitabine. Both *cds1*Δ and *rad3*Δ cells had decreased spot growth above 50 µM doses. Strangely, *cds1*Δ and *rad3*Δ had improved growth at the highest gemcitabine doses. We hypothesized that high-dose gemcitabine promotes drug resistance, particularly in *rad3*Δ cells. Large colonies also emerged at highest doses in *cds1*Δ cells after gemcitabine exposure and outgrowth. Future work will determine whether these large colonies harbour suppressor mutations caused by gemcitabine exposure. Together, our data suggest that gemcitabine effects on growth and mutation were highest in replication checkpoint mutants *rad3*Δ and *cds1*Δ, and less effective at high dose.

**Figure 3.**
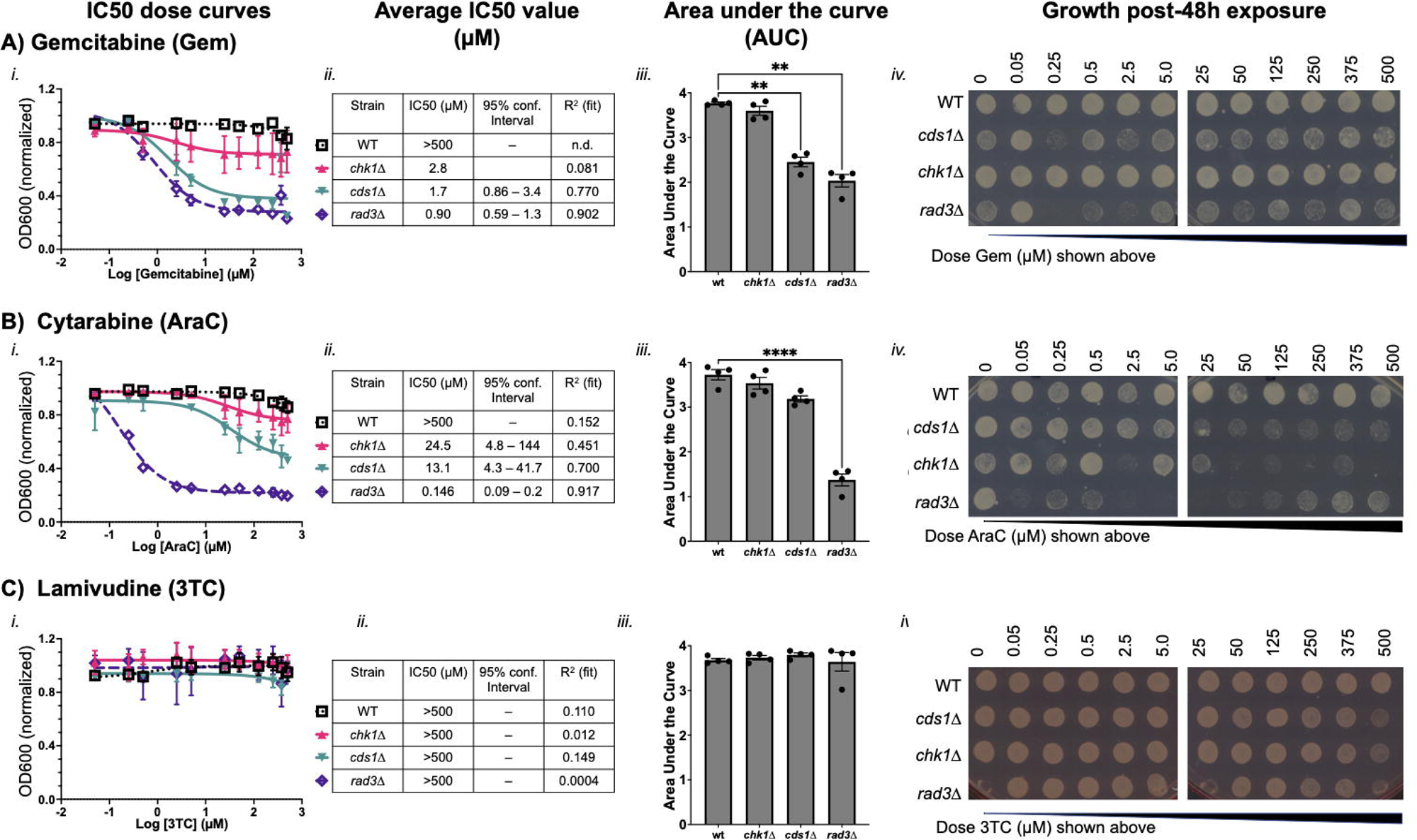
Cytidine analogues gemcitabine and AraC decrease *S. pombe* proliferation. As in Figure 2, cytidine analogues were tested in wild type (WT), chk1Δ, cds1Δ, rad3Δ strains. A) Gemcitabine (Gem), B) cytarabine (AraC), and C) lamivudine (3TC) were assessed in a minimum of 4 experimental replicates of dose response curves. An average curve of each strain/drug is presented in panel i. Standard deviation is shown around each average point. The calculated curve was used to generate an IC50 value in µM; strains that did not show growth inhibition by OD600 were described as having an IC50 above 500 µM. The area under the curve (AUC, iii) describes differences in OD600 response curves. AUC values from 4 experimental replicates were plotted and are shown with standard deviation. A one-way ANOVA was used to detect differences in AUC (** p<0.01; **** p<0.001). Absence of connecting line and asterix indicates that there was no significant difference. In iv), cells were spotted after 48h in drug onto growth media and grown in the absence of drug for 24h at 30°C. Pinned spots describe whether a strain is killed by nucleoside analogue at a given dose.

Gemcitabine did not stop culture growth in wild type cells. The wild type dose response curve, AUC values, and spot formation were all consistent with a conclusion that gemcitabine does not inhibit growth or kill cells in doses up to 500 µM. We recorded the IC50 value of gemcitabine in wild type as >500 µM from multiple replicate IC50 calculations that were averaged (Table 2). Intriguingly, when IC50 was calculated from normalized and aggregated data in the chart of Figure 3, *chk1*Δ cells generated an IC50 value of 2.8 µM gemcitabine (Figure 3Aii). Yet, the dose-response curve is not classically sigmoidal, and the degree of inhibition by gemcitabine on *chk1*Δ is low (Fig 3Aiii-lowest growth is 0.8 of normalized OD600). The area of *chk1*Δ curves in gemcitabine was not significantly different from wild type. The *chk1*Δ spots grew after all exposure doses. These data indicate that gemcitabine is not toxic to wild type or *chk1*Δ in the conditions that we tested.

**Table 2.** IC50 values calculated from independent replicates, with standard deviation. In contrast to aggregate IC50 values presented in Figures 2 and 3, this average is from non-normalized data.

| Strain | Calculated IC50 ± SD (n) |  |  |  |  |
| --- | --- | --- | --- | --- | --- |
|  | Thy | 5FU | BrdU | Gem | AraC |
| wild type (WT) | >500<sup>a</sup> (5) | 383 ± 458 (5) | 11.9 ± 4.2 (6) | >500 (4) | >500 (4) |
| *chk1Δ* | >500<sup>a</sup> (4) | >500 (4) | 7.7 ± 2.7 (4) | >500 (4) | 37.9 ± 35.8 (4) |
| *cds1Δ* | 21.6 ± 5.8 (3) | 73.6 ± 30.6 (4) | 15.9 ± 7.1 (4) | 2.2 ± 0.4 (4) | 13.6 ± 9.3 (3) |
| *rad3Δ* | 14.9 ± 1.1 (4) | 36.1 ± 16.2 (4) | 3.6 ± 1.3 (4) | 1.1 ± 0.1 (4) | 0.15 ± 0.05 (4) |

In contrast, cytarabine (AraC) decreased growth of all checkpoint mutant cultures (Figure 3B). The *rad3*Δ cells had an IC50 of 0.15 µM and a significantly lower AUC compared to wild type (IC50 AraC >500 µM). Growth of *rad3*Δ cells after AraC was tested by pinning onto plain medium, and *rad3*Δ cell growth was minimal or non-existent following even the lowest dose of 0.05 µM AraC up to 250 µM. At the 2 highest doses, *rad3*Δ cells show some growth post-exposure (Figure 3B iv). We hypothesize that high-dose AraC limits toxicity at the highest doses in *rad3*Δ. The *cds1*Δ cells had an IC50 of 13.1 µM and do not form spots above a dose of ∼5 µM. The *chk1*Δ cells had an IC50 of 24.5 µM, and do not grow above the IC50 dose of ∼25 µM. In the *cds1*Δ and *chk1*Δ samples, toxicity caused by AraC is coincident with the IC50 dose. OD600 values of non-cds1Δ cells did not capture the effect of AraC on replication-checkpoint competent cells. Replication-checkpoint mutants (*i.e. cds1*Δ) may therefore experience arrest caused by DNA damage more quickly. We propose that DNA damage induction in *cds1*Δ inhibits growth and affects OD600 and thus IC50. However, replication-checkpoint competent cells *(i.e.* wild type, *chk1*Δ) are damaged by AraC. In *chk1*Δ, the effect is lethal above 25 µM.

We tested the antiretroviral drug lamivudine (3TC, Figure 3C) to determine how a cytidine-analogue viral polymerase inhibitor was tolerated in *S. pombe* mutants. In the IC50 test, all cultures grew and an IC50 dose could not be calculated (>500 µM). AUC values were similar for all strains. However, we repeatedly saw that checkpoint mutants had reduced and slower spot growth at the highest doses of 3TC (Figure 3C iv). This spot outgrowth indicated that although 3TC is generally non-toxic, there may be some dependency on checkpoint functions to survive high 3TC doses.

### 2.3. IC50 doses of BrdU and gemcitabine induce DNA mis-segregation, whilst higher doses promote drug-resistant morphologies

During IC50 assessment, we saw that 5FU, AraC, gemcitabine, and BrdU sometimes caused more growth in spots post-treatment and above the IC50 dose. We compared drug doses between each strain over 48h, and then spotted for viability on the same plate to compare growth differences due to each drug within a strain (Figure 4A). Replication checkpoint mutant strains *cds1*Δ and *rad3*Δ showed a decrease in OD600 in nucleoside analogue around the IC50 dose. Yet, above the IC50 dose *cds1*Δ and *rad3*Δ cells often grew large and atypical colonies. Reminiscent of “suppressor” mutant colonies, these large colonies grew faster than surrounding cells on the spot background area above the IC50 dose. The highest doses frequently allowed spot growth that resembled sub-IC50 treatment. Wild type and *chk1*Δ cultures rarely showed this pattern of more growth and suppressors higher doses. Our data suggested that nucleoside analogue IC50 exposure may cause a genomic instability “crisis” in replication-checkpoint defective cells. This IC50-crisis and outgrowth presented a potential model to explore drug resistance from nucleoside analogue exposure.

**Figure 4.**
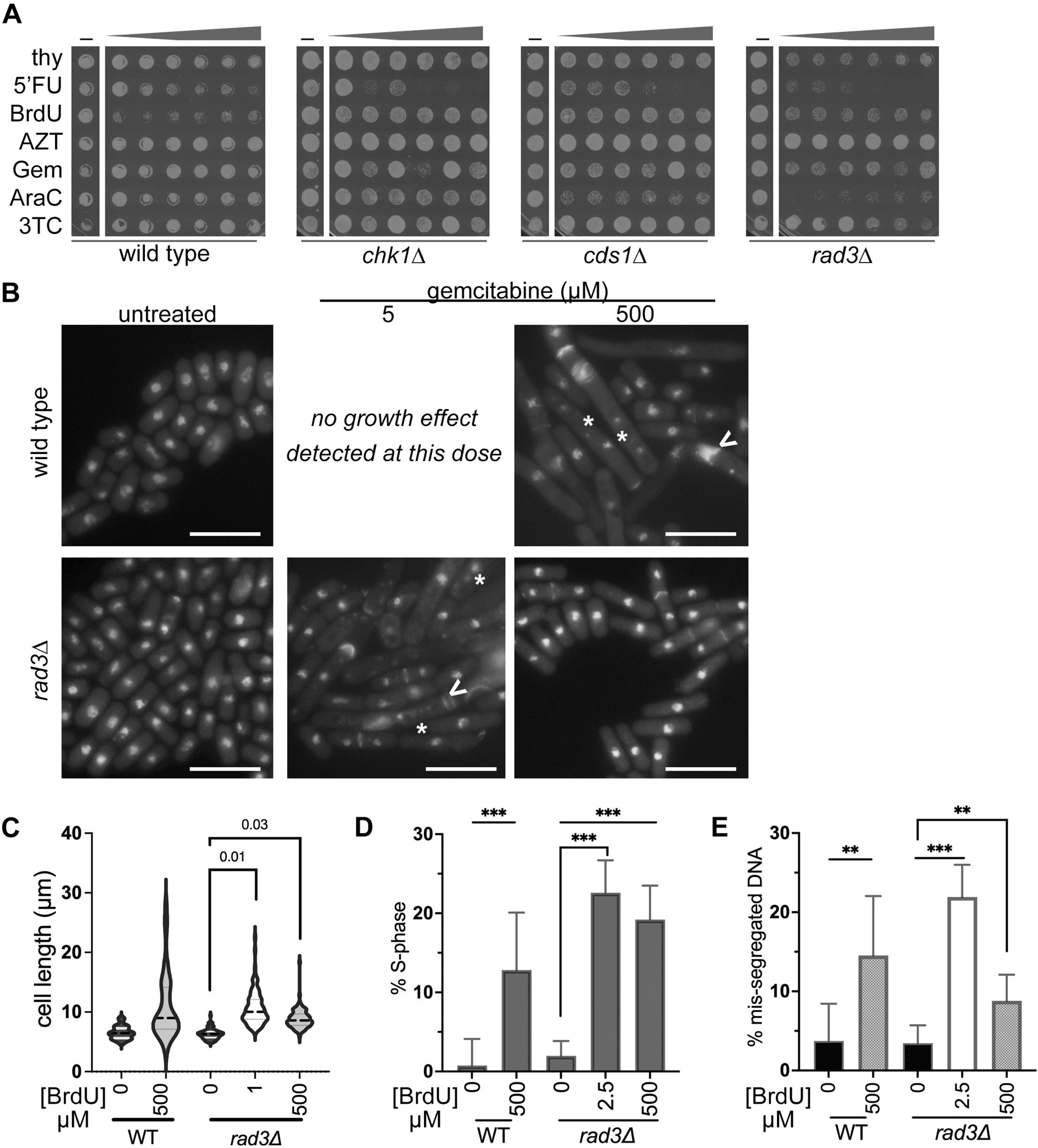
IC50 doses of BrdU and gemcitabine induce DNA mis-segregation, whilst higher doses promote drug-resistant morphologies. Cells after 48h analogue exposure were spotted onto plain YES medium and grown for 48h at 30°C. All analogues were grown and compared within the same plate to assess differences caused by each analogue, and different doses. Untreated cells are the left-most column for each strain (wild type, *chk1*Δ*, cds1*Δ*, rad3*Δ); the triangle shows increasing dose from 25 µM dose (left) to 500 µM. We found that strains that are sensitive to analogue and generate an IC50 value (Fig 2, 3) tended to generate less spot density/cell growth near the IC50 dose. *cds1*Δ and *rad3*Δ strains frequently grew better after high-dose exposure (500 µM) spots, and often generated large single colonies on small background growth (seen in this *cds1*Δ image for AraC and BrdU). B) Wild type and *rad3*Δ cells exposed to gemcitabine were fixed and stained to detect DNA and septum morphologies with no drug, IC50 (*rad3*Δ ∼ 1µM), and 500 µM. Wild type cells do not show decreased growth by OD600 and no IC50 sample was possible. Large cells, DNA mis-segregation, and septum abnormalities are seen in *rad3*Δ around the IC50 dose, like what is seen in wild type at a 500 µM dose. Scale bars 10µm. C) Cell lengths were measured and are presented in different gemcitabine doses from 3 experimental replicates. A repeated measures ANOVA was used to compare cell lengths, and calculated p values are shown above brackets. B) The *cds1*Δ cells do not show mis-segregation at the 500µM dose and have increased septation and length compared to untreated. C) Binucleate (G1) and septated (S-phase) cells were counted as a proportion of the total population from BrdU treated images. A Z-test with Yates’ continuity correction was used to compare each set (*** p<0.001). Gemcitabine causes G1/S accumulation at both doses in *rad3*Δ, and at 500 µM in wild type. ed to compare each set relative to no drug (ND). E) Gemcitabine causes mis-segregation in *rad3*Δ cells at the IC50 dose, or in wild type at 500 µM. Cells with anucleate, multiple nuclei (*), or mis-segregated DNA (<) are presented as a fraction of the total number of cells. A Z-test with Yates’ continuity correction was used to compare within each set, ** p<0.01, *** p<0.001.

Because our data suggested that the maximally effective dose for a specific mutant strain – nucleoside analogue pairing is approximately the IC50 value, we predicted that cells exposed to an IC50 dose are more likely to die and show obvious signs of genome instability. In this hypothesis, most cells in the IC50-dose treated culture die and cannot grow into spots. To compare morphology with spot results (Figure 4A), we used DNA and aniline blue septum staining [60, 61]. We imaged cells and compared morphologies under three drug conditions: no drug, the IC50 dose, and at the highest dose of 500 µM. We began with wild type and *rad3*Δ cells in gemcitabine because of the stark difference in

IC50 response. Wild type and *chk1*Δ cultures did not generate a sigmoidal curve or IC50 value in gemcitabine using OD600; we inferred that DNA damage response is not essential to stopping growth. Because wild type cells were insensitive to gemcitabine and did not generate an IC50 value, lower-dose IC50-dose cells could be tested.

However, this did not mean that wild type cells were unaffected by gemcitabine treatment. We found that wild type cells in 500 µM gemcitabine were long and multi-septated, with clear DNA mis-segregation (Figure 4B). We compared wild type cell length in untreated and 500 µM gemcitabine and found that gemcitabine treatment caused cell elongation (Figure 4C). The number of anaphase and septated wild type cells was higher in 500 µM gemcitabine, showing that G1-S phase arrest occurs (Figure 4D). Nuclear mis-segregation was increased in wild type after 500 µM gemcitabine (Figure 4E). We conclude that wild type cells are sensitive to high-dose gemcitabine (Figure 3Ai, black line) reflects gemcitabine sensitivity, even though no lower-dose IC50 is calculated. Further, we infer that wild type DNA damage checkpoint prevents division in gemcitabine, correlated with cell elongation and septation phenotypes.

In contrast, *rad3*Δ cells had a gemcitabine IC50 value of approximately 1 µM (Table 2). *rad3*Δ cells at the IC50 dose showed multiple nuclei, *cut* DNA, elongation, and multiple septa (Figure 4B). This was consistent with IC50 doses of gemcitabine causing genomic instability in *rad3*Δ cells. At the highest 500 µM dose of gemcitabine, *rad3*Δ cells were more uniform in size, although longer than untreated (Figure 4C, Table 3); we attribute some of this difference to cell growth arrest in untreated cultures after 48h static growth. More *rad3*Δ cells had septa or anaphase nuclei in either IC50 or 500 µM gemcitabine, indicating more cells were in G1-S phases (Figure 4E). Morphologically, *rad3*Δ cells in IC50 gemcitabine looked like wild type cells at the highest (500 µM) dose (Figure 4B). By counting abnormal nuclear segregation of DAPI, we found that *rad3*Δ cells at 500 µM gemcitabine had fewer obvious DNA mis-segregation events than at the IC50 dose (Figure 4D). Our data indicate that *rad3*Δ cells are most sensitive to gemcitabine at the IC50 dose where genome instability is obvious.

**Table 3.** Cells are longest in gemcitabine concentration near the IC50 dose for *rad3*Δ. The *rad3*Δ cells are affected by either 1 µm or 500 µM gemcitabine. Despite no obvious gemcitabine sensitivity, wild type cells are longer at 500 µM gemcitabine dose. Tip-to-tip or tip-to-septum mean cell lengths are shown in µm +/- standard deviation. The p value indicates if there is a statistically significant difference between the cell lengths at each dose of drug relative to no treatment. A repeated measures one-way ANOVA test was used, assuming Gaussian distribution and the Geisser-Greenhouse correction for matched data sets. Tukey’s multiple comparisons correction was applied.

|  | Gem dose (μM) | Mean length +/-SD (μm) | Number of cells (n) | P value (untreated compared to treated) |
| --- | --- | --- | --- | --- |
| Wild type | 0 | 6.5 (1.1) | 100 | — |
|  | 500 | 10.9 (5.5) | 105 | 0.005 |
| *rad3Δ* | 0 | 6.3 (0.1) | 301 (3) | — |
|  | IC50 | 10.7 (0.7) | 312 (3) | 0.01 |
|  | 500 | 8.8 (0.5) | 304 (3) | 0.03 |

Because 500 µM gemcitabine doses impacted *rad3*Δ cells less, we hypothesized that other nucleoside analogues may arrest replication checkpoint mutant cells, but not kill at 500 µM doses. We tested BrdU, because all strains were sensitive to BrdU and we calculated IC50 doses (Figure 2C, Table 2). We predicted that *cds1*Δ or *rad3*Δ cells at an IC50 dose would show more DNA mis-segregation and abnormal morphologies (*i.e*. longer cells or septated/anaphase cell accumulation) than at the highest dose. Instead, we found that all genotypes showed dramatic changes in morphology at their respective BrdU IC50 doses (Figure 5A), including elongation, abnormal septation, aneuploidy, and multinucleated cells. Intriguingly, cells that were treated with 500 µM BrdU had morphologies like untreated and cycling fission yeast.

**Figure 5.**
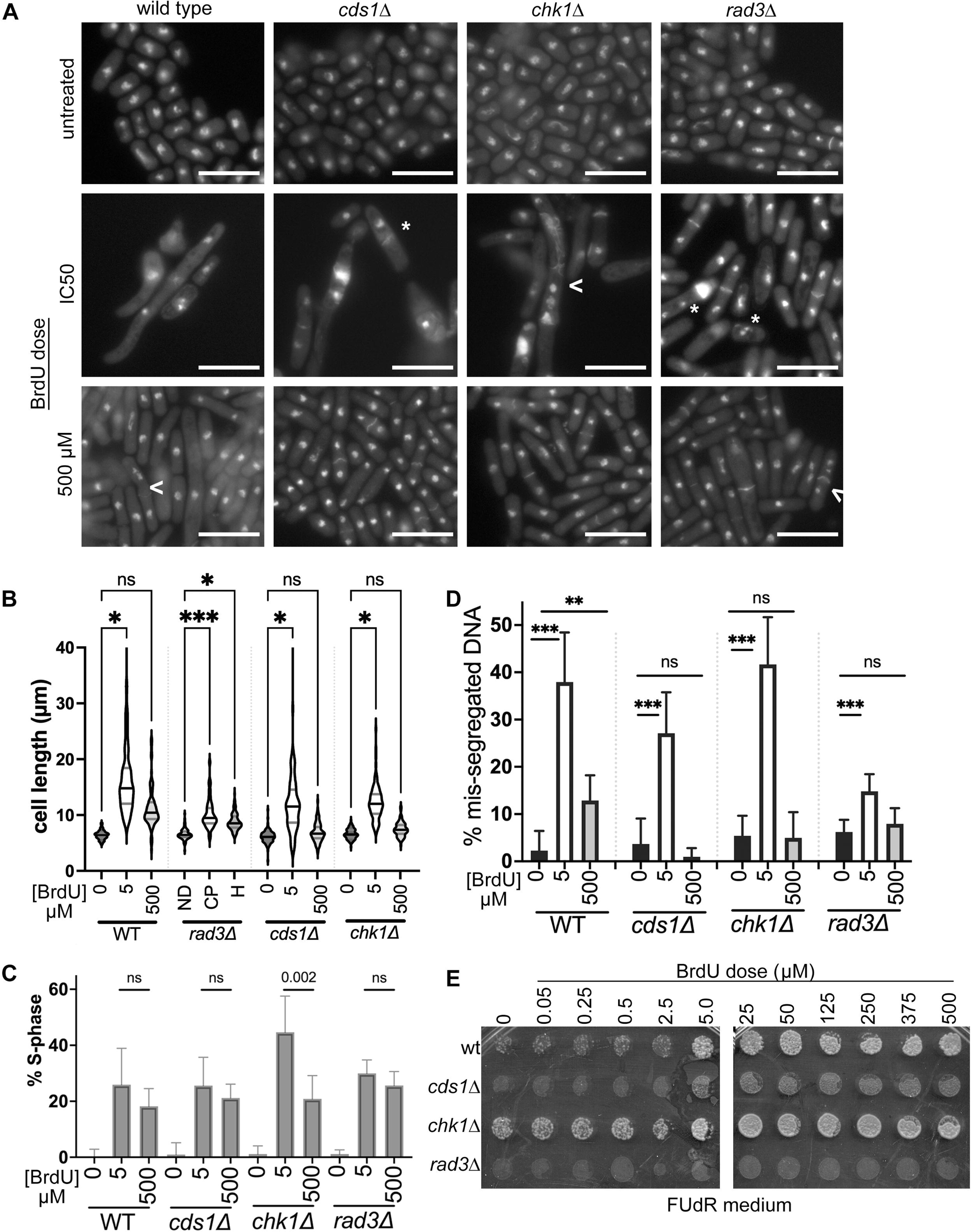
IC50 doses of BrdU cause elongation, G1/S accumulation, and DNA mis-segregation that alter nucleoside analogue resistance. A) Microscopy images were assessed for cell elongation as in Figure 4 (scale bars 10 µm) at 48h of BrdU treatment. Frequent DNA mis-segregation events including anaphase and additional nuclei (*) or septation errors (<) were observed in all cells at the IC50 dose of 5-25 µM BrdU. B) Tip-to-tip cell length increases in BrdU and is longest at the IC50 dose. Three (3) experimental replicates were assessed using a violin plot with mean and 25% / 75% quartiles (note-population averages and SD reported in Table 4). The difference between no BrdU and treatments in each strain was calculated using a repeated measures ANOVA with correction (* p<0.05; *** p<0.001, ns not significant). C) Binucleate (G1) and septated (S-phase) cells were counted as a proportion of the total population in BrdU treated images. A Z-test with Yates’ continuity correction was used to compare each set. Each strain has a statistically significant increase in G1/S-% with either BrdU treatment compared to no BrdU. In chk1Δ, the proportion of G1/S cells is increased at the 5 µM IC50 dose compared to 500 µM. D) Nuclear mis-segregation is increased in BrdU at an IC50 dose (5 µM) in all strains. Cells with atypical DNA segregation, including anucleate, cut, multiple nuclei, or lagging chromosomes, were counted as a proportion of the total number of cells. A Z-test with Yates’ continuity correction was used to compare within each strain, ns – not significant, ** p<0.01, *** p<0.001. Wild type cells show more mis-segregation at 500 µM compared to no BrdU. All other strains did not have a statistically different proportion of mis-segregation at 500 µM. E) After 48 h BrdU exposure, cultures were spotted onto FUdR to test for hsv-tk^+^ function. Nucleoside analogue sensitive cells require hsv-tk^+^ for sensitivity to BrdU (0 µM dose column). With increasing BrdU, spots acquire additional FUdR-insensitive cells, even as the number of viable cells decreases (compare to Fig 2Civ).

**Table 4.** All strains elongate in BrdU at an IC50 dose. Cell lengths were measured without BrdU (0 µM), in an IC50 dose (used 5 µM as an average of IC50 values), or in 500 µM BrdU. Cells were measured tip-to-tip, or septum to tip. Three (3) experimental populations were measured, and N used in statistics was 3; the total number of cells within these replicates is reported (n). Average cell length is compared between dose for each strain. A repeated measure one-way ANOVA test was used with Tukey’s correction. Each p-value indicates whether the difference between untreated or BrdU treatment is significantly different (* p<0.01; *** p<0.001; not significant, ns).

| BrdU | BrdU dose (μM) | Mean length (+/-SD, μm) | Number of cells (n) | P value (significance) |
| --- | --- | --- | --- | --- |
| Wild type | 0 | 6.5 (0.9) | 289 | — |
|  | 5 | 15.9 (1.9) | 317 | 0.05 (*) |
|  | 500 | 11.1 (1.1) | 282 | 0.10 (ns) |
| *cds1Δ* | 0 | 6.3 (0.4) | 282 | — |
|  | 5 | 12.5 (2.1) | 224 | 0.05 (*) |
|  | 500 | 6.9 (0.3) | 208 | 0.12 (ns) |
| *chk1Δ* | 0 | 6.7 (0.5) | 287 | — |
|  | 5 | 12.3 (0.7) | 269 | 0.02 (*) |
|  | 500 | 7.6 (1.3) | 306 | 0.60 (ns) |
| *rad3Δ* | 0 | 6.5 (0.4) | 306 | — |
|  | 5 | 10.1 (0.3) | 292 | <0.0001 (***) |
|  | 500 | 8.7 (0.3) | 308 | <0.04 (*) |

*S. pombe* cells elongate during G2 arrest and in response to DNA damage and may show increased DNA mis-segregation as a symptom of chromosome instability. Elongation could also contribute to higher OD600 measurements which might affect growth curves. To test for elongation, we measured cell lengths after IC50 and high-dose BrdU. We found that BrdU at low- or high-dose caused longer cells and more variation (Figure 5B). Wild type cells elongated in IC50 doses and remained longer at high dose (Table 4); this implies that there was a G2-arrest or block to cytokinesis. In contrast, checkpoint mutants showed longer cells at the IC50 dose, the populations were generally shorter at 500 µM (Table 4). BrdU increased the proportion of binucleate and septated cells at either dose (Figure 5C), indicating G1/S accumulation and arrest. DNA mis-segregation increased in all cells and at either IC50 or 500 µM doses. However, the IC50 dose of BrdU caused the highest frequency of mis-segregation in all cells (Figure 5D), showing that the dose of most sensitivity is correlated with the largest biological effect. Checkpoint mutants had less mis-segregation at 500 µM BrdU, with no significant difference between 500 µM and untreated.

Finally, we tested whether cells were still sensitive to nucleoside analogue treatment after BrdU (Figure 5E, Supplementary Figure S1). We spotted cultures onto 5-fluoro-2’-deoxyuridine (FUdR) to assess HSV-TK function. FUdR added to medium detects the emergence of nucleoside analogue-resistant cells, because analogue can no longer be phosphorylated [6, 7]. We saw that wild type, *cds1*Δ, and *rad3*Δ cells had few FUdR-resistant cells after incubation without BrdU. In contrast, *chk1*Δ cells have a higher frequency of FUdR-resistance that may indicate general genome instability and HSV-TK loss even without BrdU. With increased BrdU dose the number of FUdR-resistant colonies within each spot increased. This occurred even as the overall number of cells per dot decreased with increasing FUdR (Figure 2Civ). We spotted onto canavanine to assess forward mutation of canavanine sensitivity. We found that *cds1*Δ and *chk1*Δ cells have more FUdR growth in lower-concentration spots, up to the IC50. Contrastingly, the *rad3*Δ cells have FUdR-resistant colonies that form in doses up to the IC50, and again at the highest doses of BrdU (Supplementary Figure S1).

We conclude that lower-dose BrdU promotes DNA replication arrest and DNA damage induction. An IC50 BrdU dose is apparently more toxic to cells. However, high-dose BrdU above the IC50 dose generates a different spectrum of damage and/or mutation. Based on our FUdR tests, the changes caused by BrdU may decrease nucleoside analogue sensitivity by decreased HSV-TK function; this may promote cell survival and drug resistance. Because lower DNA mis-segregation was seen in the at highest doses of BrdU (Figure 5D), we propose that high-dose BrdU induces changes that promote survival, decrease nucleoside analogue toxic effects, and may promote downstream drug resistance.

## 3. Conclusions

Using a rapid synthetic biology method for *S. pombe*, we show that nucleoside analogue drug dose outcome is influenced by cellular DNA replication (Cds1-dependent) or DSB repair (Chk1-dependent) checkpoints, consistent with previous literature in other models and methods [50, 62–64]. Thus, fission yeast liquid growth screens using OD600 can identify drug mechanisms [49] and mutational outcomes [64]. Capitalizing on the similarities of *S. pombe* and human checkpoint proteins and dNTP metabolism pathways [6, 45], we show that the IC50 dose for cytotoxic nucleoside analogues is linked to checkpoint competency. IC50 is lowest with loss of the ATR-homologue Rad3. Drug mechanisms specific to each analogue may influence IC50 dose, treatment failure, or risk of resistance. An IC50 dose of nucleoside analogue causes DNA mis-segregation and G1/S-phase arrest. The relationship between genetic mutation background and IC50 dose of required to reduce cell growth (Figures 2, 3; Table 2) maximizes cell killing without off-target effects.

However, BrdU and gemcitabine doses above a strain-specific IC50 dose allowed growth recovery and decreased FUdR sensitivity (Figures 4, 5). FUdR insensitivity implies that HSV-TK function was reduced. Doses above the IC50 dose also caused large colonies suggestive of suppressor mutations, and fewer morphological changes in cell length or DNA mis-segregation. Thus, nucleoside analogue doses that are too high may increase unanticipated drug insensitivity and risk of later resistance. Because gemcitabine sensitivity depends on checkpoint competency and growth medium [49, 50, 64], failing to consider genomics and outcome relative to dose may lead to incorrect dosing, drug insensitivity, and mutational changes within a once-sensitive population of cells.

A predictable relationship between a specific nucleoside analogue, an individual’s cellular genotype or phenotype, drug sensitivity range, and loss of sensitivity threshold may allow decreased drug use with synthetic lethality principles. Synthetic lethality uses 2 conditions that are non-lethal independently to cause better cell killing when together [65, 66]. In cancer, a synthetic lethality strategy could target tumour cells for nucleoside analogue effects, maximizing tumour death, minimizing risk of resistance, and sparing non-cancerous “bystander” cells. Our method can be applied to find a specific IC50 dose that reduces cell growth (Figures 2, 3; Table 2), while minimizing the risk of failure or resistance above the IC50. Testing for suppressor mutations that develop at the IC50 dose in checkpoint mutant cells (i.e. *cds1*Δ*, rad3*Δ) may improve treatments and predict causes of drug resistance.

However, our work shows a caution for ATR inhibition with nucleoside analogues: because ATR is the homologue of *S*. *pombe* Rad3 and we see off-target effects in *rad3*Δ above the IC50 dose and risk of resistance, the use of ATR inhibitors to increase nucleoside analogue effects (*i.e*. [67, 68]) could confer a higher risk of mutation and later drug resistance.

Patient paired genotype-dose targeting supports patient quality of life during chronic HIV prophylaxis and cancer therapy. Because nucleoside analogues target proliferating cells such as hair follicles, intestinal epithelium, and bone marrow, drug side effects can include hair loss, nausea, anemia, and compromised immunity. Even antiretroviral drugs such as AZT and 3TC can cause these effects, and also lipodystrophy and metabolic diseases (e.g. hyperglycemia, hypercholesterolemia)[69, 70]. AZT and 3TC do not typically target DNA polymerase, kill uninfected cells, or stop proliferation; our data that higher 3TC doses change *rad3*Δ spot growth supports effects that are subtle and more mutation-based over long-term treatment [71]. Patient side-effects can cause therapy refusal leading to co-morbidities, disease progression, and accelerated drains on patient-caregiver-health networks, even if the drug is effective [72]. Using the lowest possible dose improves quality of life and may extend the window of specific nucleoside analogue use before a switch is required due to acquired drug resistance.

We conclude that personal genomics profiling ahead of treatment may predict outcomes. *S. pombe* screening using these methods could be used to amplify nucleoside analogue sensitivities and minimize the risk of resistance during long-term use. The *S. pombe wee1*Δ mutant led to the discovery of Wee1 inhibitor Adavosertib. Adavosertib is now in clinical trials targeting uterine, lung, ovarian, renal and refractory tumours [59, 73, 74]. Wee1 inhibition can synergistically enhance radiation and gemcitabine treatment in cancer [73, 75–77]. Thus, targeting inhibitors of cell cycle and checkpoint kinases may improve nucleoside analogue drug use through synthetic lethality, while avoiding mutagenesis and risk of later insensitivity.

## 4. Materials and Methods

### 4.1. Yeast strains

Fission yeast cells were cultured following standard conditions *e.g.*[78, 79]. Cells were grown from frozen stocks on yeast extract with supplements (adenine, histidine, leucine, uracil, lysine; YES 225 formulation, Sunrise Bioscience, CA, USA) at 30°C. Cultures for nucleoside analogue exposure were incubated at 30°C overnight in Pombe Minimal Glutamate (PMG) media supplemented with uracil and adenine (both 225 mg/L; BioShop, Canada). Other supplements were added as appropriate at 225 mg/L. Strain genotypes are listed in Table 1.

*S. pombe* lacks a thymidine salvage pathway. Strains used to assess nucleoside analogue sensitivity contained Herpes Simplex Virus 1 (HSV1) thymidine kinase that was stably integrated and expressed from a constitutive alcohol dehydrogenase promoter (*Padh1*-*hsv-tk^+^).* The human equilibrative nucleoside transporter 1 (hENT1) was stably integrated into strains using the *Padh1* promoter (*Padh1*-*hENT1^+^).* Expression of hENT1 allows better nucleoside analogue uptake at lower doses as described in [6, 52].

### 4.2. Drugs

Nucleoside analogue drugs were resuspended in water or DMSO as stock solutions of 100 µM. The drugs used include gemcitabine (Toronto Research Chemicals, Canada), cytarabine (AraC; SigmaAldrich Canada), 5-fluorouracil (5-FU; BioShop Canada), 5-fluoro-2’-deoxyuridine (FUdR; BioShop Canada), bromodeoxyuridine (BrdU; SigmaAldrich Canada), thymidine (BioShop, Canada). Anti-retroviral compounds zidovudine (AZT) and lamivudine (3TC) were kind donations from Dr Russell Viirre (Toronto Metroolitan University, Canada). Canavanine sulfate (Cedarlane, Canada) was added to PMG+HULA medium in Supplementary Figure 1 at 70 µg/mL [80].

### 4.3. Proliferation assay and IC50

Liquid culture growth was used to evaluate the dependency of nucleoside analogue response on the cell cycle checkpoint. Decreased proliferation in drug could indicate cell death or arrest. *S. pombe* checkpoint mutant cells (Table 1) were grown overnight in pombe minimal glutamate (PMG) media supplemented with 225 mg/mL uracil and adenine. Cultures were diluted in fresh PMG + UA to an OD600 of approximately 0.1. Diluted cultures were kept at room temperature for 2 hours to acclimate. Well plates were prepared containing 200 µL of diluted cells, PMG+UA medium, and drug. Each drug was prepared and used at final concentrations of 0, 0.05, 0.25, 0.5, 2.5, 5.0, 25, 50, 125, 250, 375, 500 µM; these doses generate an 11-point IC50 curve.

Optical density measurements at a wavelength of 600 nm (OD600) were taken using a spectrophotometer in 96-well growth plates. OD600 readings were taken before static incubation at 30°C, and after 24h and 48h of growth at 30°C. After the 48h OD600 reading, 96 well plates were spotted onto yeast extract with supplements (YES; Sunrise Media, Carlsbad, CA USA) solid media (1.8% agar; BioShop) prepared and autoclaved as described. Spotting was performed using a 48-array pinner (V&P Scientific, 3 µL hanging droplet). Pinned plates were then incubated at 30°C overnight. The following day, the YES plates were scanned for analysis. In supplementary figure S1 and Figure 5, cells were also spotted onto PMG + HULA containing either FUdR or 70 µg/mL canavanine with Phloxine B. Spots were incubated 3 days (FUdR) or 9 days (canavanine) before assessment.

### 4.4. Microscopy

Cells were fixed in cold 100% ethanol to a final concentration of 70% ethanol. Samples were vortexed and stored at 4°C. To stain, fixed cells were rehydrated in water, and then incubated in aniline methylene blue solution (AMB; 1% aniline blue water-soluble powder w/v, in 1xPBS) for 15 minutes at room temperature. AMB binds to the *S. pombe* septum and fluoresces to specifically reveal septa. AMB-stained cells were smeared onto glass slides and air dried at room temperature. Samples were mounted in 4 μL of fluorescent imaging mounting medium (50% glycerol, 1% DABCO) containing 1 μg/mL DAPI. Samples were sealed with a coverslip and stored in a light-protected slide box at −20°C. Cells were imaged on an Olympus IX83 Inverted Microscope linked to a Hamamatsu ORCA-Flash 4.0 CMOS digital camera. CellSens acquisition software (Olympus) was used to acquire images on DAPI-excitation 359 nm/emission 461 nm, and bright field settings. FIJI software was used to analyse the images [81].

### 4.5. Statistical analysis

IC50 OD600 values were imported into GraphPad Prism 10 for analysis. Drug concentrations were log10-transformed. A 3-parameter, non-linear fit of dose-response was fit to the data. IC50 values were calculated for each independent replicate. Graphs were made using the 48h OD600 readings for comparison with calculated IC50 values. Replicate IC50 values were averaged and assessed using standard deviation between calculated values. Microscopy images were examined, and cells were measured tip-to-tip or tip-to-septum. A pairwise two-tailed t-test was used to compare cell lengths between treatment conditions and genotypes (GraphPad Prism 10).

## Author Contributions

Conceptualization, Sarah Sabatinos; Data curation, Sarah Sabatinos; Formal analysis, Zainab Kagalwala, Mohammed Ayan Chhipa and Sarah Sabatinos; Funding acquisition, Sarah Sabatinos; Investigation, Zainab Kagalwala, Mohammed Ayan Chhipa and Zohreh Kianfard; Methodology, Zainab Kagalwala, Mohammed Ayan Chhipa, Sirasie Magalage, Zohreh Kianfard, Essam Karam and Sarah Sabatinos; Project administration, Sarah Sabatinos; Supervision, Sarah Sabatinos; Writing – original draft, Zainab Kagalwala and Sarah Sabatinos; Writing – review & editing, Mohammed Ayan Chhipa, Zohreh Kianfard, Essam Karam, Sirasie Magalage and Sarah Sabatinos.

## Institutional Review Board Statement

Not applicable.

## Informed Consent Statement

Not applicable.

## Data Availability Statement

Data supporting reported results are available upon request of Sarah Sabatinos.

## Acknowledgments

We thank Ella Hyatt and Salwa Saeed at Toronto Metropolitan University for their administrative and facility support that was critical to research productivity. We thank the University College London Pharmacy Program for their support and program/student support. We thank members of the Sabatinos Lab for critical reading of this manuscript.

## REFERENCES

1. Galmarini CM, Mackey JR, Dumontet C (2002) Nucleoside analogues and nucleobases in cancer treatment. Lancet Oncol 3:415–424

2. Menéndez-Arias L (2008) Mechanisms of resistance to nucleoside analogue inhibitors of HIV-1 reverse transcriptase. Virus Res 134:124–146. 10.1016/j.virusres.2007.12.015

3. Tsesmetzis N, Paulin CBJ, Rudd SG, Herold N (2018) Nucleobase and Nucleoside Analogues: Resistance and Re-Sensitisation at the Level of Pharmacokinetics, Pharmacodynamics and Metabolism. Cancers 10:240. 10.3390/cancers10070240

4. Peters GJ (2014) Novel Developments in the Use of Antimetabolites. Nucleosides Nucleotides Nucleic Acids 33:358–374. 10.1080/15257770.2014.894197

5. Mead TJ, Lefebvre V (2014) Proliferation Assays (BrdU and EdU) on Skeletal Tissue Sections. In: Hilton MJ (ed) Skeletal Development and Repair. Humana Press, Totowa, NJ, pp 233–243

6. Sabatinos SA, Mastro TL, Green MD, Forsburg SL (2013) A mammalian-like DNA damage response of fission yeast to nucleoside analogs. Genetics 193:143–57. 10.1534/genetics.112.145730

7. Sivakumar S, Porter-Goff M, Patel PK, et al (2004) In vivo labeling of fission yeast DNA with thymidine and thymidine analogs. Methods 33:213–9. 10.1016/j.ymeth.2003.11.016

8. (2010) Zidovudine and Lamivudine for HIV Infection. Clin Med Rev Ther 2:115–127. 10.4137/CMRT.S4557

9. Weinert T (1998) DNA damage and checkpoint pathways: molecular anatomy and interactions with repair. Cell 94:555–558

10. Alcasabas AA, Osborn AJ, Bachant J, et al (2001) Mrc1 transduces signals of DNA replication stress to activate Rad53. Nat Cell Biol 3:958–65. 10.1038/ncb1101-958

11. Enoch T, Carr AM, Nurse P (1992) Fission yeast genes involved in coupling mitosis to completion of DNA replication. Genes Dev 6:2035–2046. 10.1101/gad.6.11.2035

12. Lindsay HD, Griffiths DJ, Edwards RJ, et al (1998) S-phase-specific activation of Cds1 kinase defines a subpathway of the checkpoint response in Schizosaccharomyces pombe. Genes Dev 12:382– 95

13. Boddy MN, Furnari B, Mondesert O, Russell P (1998) Replication checkpoint enforced by kinases Cds1 and Chk1. Science 280:909–12

14. Brondello JM, Boddy MN, Furnari B, Russell P (1999) Basis for the checkpoint signal specificity that regulates Chk1 and Cds1 protein kinases. Mol Cell Biol 19:4262–9

15. Zeng Y, Forbes KC, Wu Z, et al (1998) Replication checkpoint requires phosphorylation of the phosphatase Cdc25 by Cds1 or Chk1. Nature 395:507–10. 10.1038/26766

16. Crasta K, Ganem NJ, Dagher R, et al (2012) DNA breaks and chromosome pulverization from errors in mitosis. Nature 482:53–8. 10.1038/nature10802

17. Zhang CZ, Spektor A, Cornils H, et al (2015) Chromothripsis from DNA damage in micronuclei. Nature 522:179–84. 10.1038/nature14493

18. Zhang F, Carvalho CM, Lupski JR (2009) Complex human chromosomal and genomic rearrangements. Trends Genet 25:298–307

19. Holland AJ, Cleveland DW (2012) Chromoanagenesis and cancer: mechanisms and consequences of localized, complex chromosomal rearrangements. Nat Med 18:1630–8. 10.1038/nm.2988

20. Shoshani O, Bakker B, de Haan L, et al (2021) Transient genomic instability drives tumorigenesis through accelerated clonal evolution. Genes Dev 35:1093–1108. 10.1101/gad.348319.121

21. Garcia-Closas M, Hall P, Nevanlinna H, et al (2008) Heterogeneity of breast cancer associations with five susceptibility loci by clinical and pathological characteristics. PLoS Genet 4:e1000054. 10.1371/journal.pgen.1000054

22. Vargas-Rondón N, Villegas VE, Rondón-Lagos M (2017) The Role of Chromosomal Instability in Cancer and Therapeutic Responses. Cancers 10:. 10.3390/cancers10010004

23. Lepage CC, Morden CR, Palmer MCL, et al (2019) Detecting Chromosome Instability in Cancer: Approaches to Resolve Cell-to-Cell Heterogeneity. Cancers 11:. 10.3390/cancers11020226

24. Sahin IH, Lowery MA, Stadler ZK, et al (2016) Genomic instability in pancreatic adenocarcinoma: a new step towards precision medicine and novel therapeutic approaches. Expert Rev Gastroenterol Hepatol 10:893–905. 10.1586/17474124.2016.1153424

25. Mansoori B, Mohammadi A, Davudian S, et al (2017) The Different Mechanisms of Cancer Drug Resistance: A Brief Review. Adv Pharm Bull 7:339–348. 10.15171/apb.2017.041

26. Wu L, Tannock IF (2003) Repopulation in murine breast tumors during and after sequential treatments with cyclophosphamide and 5-fluorouracil. Cancer Res 63:2134–8

27. Davis AJ, Tannock IF (2002) Tumor physiology and resistance to chemotherapy: repopulation and drug penetration. Cancer Treat Res 112:1–26

28. Vukovic B, Beheshti B, Park P, et al (2007) Correlating breakage-fusion-bridge events with the overall chromosomal instability and in vitro karyotype evolution in prostate cancer. Cytogenet Genome Res 116:1–11. 10.1159/000097411

29. Bianchi V, Pontis E, Reichard P (1986) Changes of deoxyribonucleoside triphosphate pools induced by hydroxyurea and their relation to DNA synthesis. J Biol Chem 261:16037–42

30. Alvino GM, Collingwood D, Murphy JM, et al (2007) Replication in hydroxyurea: it’s a matter of time. Mol Cell Biol 27:6396–406. 10.1128/MCB.00719-07

31. Errico A, Costanzo V, Hunt T (2007) Tipin is required for stalled replication forks to resume DNA replication after removal of aphidicolin in Xenopus egg extracts. Proc Natl Acad Sci U A 104:14929–34

32. Kurose A, Tanaka T, Huang X, et al (2006) Effects of hydroxyurea and aphidicolin on phosphorylation of ataxia telangiectasia mutated on Ser 1981 and histone H2AX on Ser 139 in relation to cell cycle phase and induction of apoptosis. Cytom A 69:212–21

33. Seifer M, Hamatake RK, Colonno RJ, Standring DN (1998) In vitro inhibition of hepadnavirus polymerases by the triphosphates of BMS-200475 and lobucavir. Antimicrob Agents Chemother 42:3200–8

34. Rinaldi C, Pizzul P, Longhese MP, Bonetti D (2021) Sensing R-Loop-Associated DNA Damage to Safeguard Genome Stability. Front Cell Dev Biol 8:618157. 10.3389/fcell.2020.618157

35. Promonet A, Padioleau I, Liu Y, et al (2020) Topoisomerase 1 prevents replication stress at R-loop-enriched transcription termination sites. Nat Commun 11:3940. 10.1038/s41467-020-17858-2

36. Zimmer J, Tacconi EM, Folio C, et al (2016) Targeting BRCA1 and BRCA2 Deficiencies with G-Quadruplex-Interacting Compounds. Mol Cell 61:449–60. 10.1016/j.molcel.2015.12.004

37. Chang EY-C, Tsai S, Aristizabal MJ, et al (2019) MRE11-RAD50-NBS1 promotes Fanconi Anemia R-loop suppression at transcription-replication conflicts. Nat Commun 10:4265. 10.1038/s41467-019-12271-w

38. Lisby M, Barlow JH, Burgess RC, Rothstein R (2004) Choreography of the DNA damage response: spatiotemporal relationships among checkpoint and repair proteins. Cell 118:699–713

39. Du LL, Nakamura TM, Russell P (2006) Histone modification-dependent and -independent pathways for recruitment of checkpoint protein Crb2 to double-strand breaks. Genes Dev 20:1583–96. 10.1101/gad.1422606

40. Jazayeri A, Falck J, Lukas C, et al (2006) ATM- and cell cycle-dependent regulation of ATR in response to DNA double-strand breaks. Nat Cell Biol 8:37–45. 10.1038/ncb1337

41. Hicks WM, Yamaguchi M, Haber JE (2011) Real-time analysis of double-strand DNA break repair by homologous recombination. Proc Natl Acad Sci U S A 108:3108–15. 10.1073/pnas.1019660108

42. Mehta A, Haber JE (2014) Sources of DNA double-strand breaks and models of recombinational DNA repair. Cold Spring Harb Perspect Biol 6:a016428. 10.1101/cshperspect.a016428

43. Acilan C, Potter DM, Saunders WS (2007) DNA repair pathways involved in anaphase bridge formation. Genes Chromosomes Cancer 46:522–31. 10.1002/gcc.20425

44. Liu Y, Nielsen CF, Yao Q, Hickson ID (2014) The origins and processing of ultra fine anaphase DNA bridges. Curr Opin Genet Dev 26:1–5. 10.1016/j.gde.2014.03.003

45. Hoffman CS, Wood V, Fantes PA (2015) An Ancient Yeast for Young Geneticists: A Primer on the Schizosaccharomyces pombe Model System. Genetics 201:403–423. 10.1534/genetics.115.181503

46. Sipiczki M (2000) Where does fission yeast sit on the tree of life? Genome Biol 1:reviews1011.1. 10.1186/gb-2000-1-2-reviews1011

47. Vyas A, Freitas AV, Ralston ZA, Tang Z (2021) Fission Yeast Schizosaccharomyces pombe: A Unicellular “Micromammal” Model Organism. Curr Protoc 1:e151. 10.1002/cpz1.151

48. Chhipa MA, Sanayhie SA, Sabatinos SA (2025) Assessing Drug Sensitivity in Fission Yeast Using Half-Maximal Inhibitory Concentration (IC50) Assays. Methods Mol Biol Clifton NJ 2862:241–253. 10.1007/978-1-0716-4168-2_17

49. Boeckemeier L, Kraehenbuehl R, Keszthelyi A, et al (2020) Mre11 exonuclease activity removes the chain-terminating nucleoside analog gemcitabine from the nascent strand during DNA replication. Sci Adv 6:eaaz4126. 10.1126/sciadv.aaz4126

50. Alyahya MY, Khan S, Bhadra S, et al (2022) Replication stress induced by the ribonucleotide reductase inhibitor guanazole, triapine and gemcitabine in fission yeast. FEMS Yeast Res 22:foac014. 10.1093/femsyr/foac014

51. Fleck O, Fahnøe U, Løvschal K, et al (2017) Deoxynucleoside Salvage in Fission Yeast Allows Rescue of Ribonucleotide Reductase Deficiency but Not Spd1-Mediated Inhibition of Replication. Genes 8:128. 10.3390/genes8050128

52. Hodson JA, Bailis JM, Forsburg SL (2003) Efficient labeling of fission yeast Schizosaccharomyces pombe with thymidine and BUdR. Nucleic Acids Res 31:e134

53. Rhind N, Russell P (2000) Chk1 and Cds1: linchpins of the DNA damage and replication checkpoint pathways. J Cell Sci 113 ( Pt 22):3889–96

54. Bentley NJ, Holtzman DA, Flaggs G, et al (1996) The Schizosaccharomyces pombe rad3 checkpoint gene. EMBO J 15:6641–6651

55. Sabatinos SA, Green MD, Forsburg SL (2012) Continued DNA synthesis in replication checkpoint mutants leads to fork collapse. Mol Cell Biol 32:4986–97. 10.1128/MCB.01060-12

56. Yanagida M (1998) Fission yeast cut mutations revisited: control of anaphase. Trends Cell Biol 8:144–149. 10.1016/s0962-8924(98)01236-7

57. Chen G, Deng X (2018) Cell Synchronization by Double Thymidine Block. Bio-Protoc 8:e2994. 10.21769/BioProtoc.2994

58. Hung C-W, Martínez-Márquez JY, Javed FT, Duncan MC (2018) A simple and inexpensive quantitative technique for determining chemical sensitivity in Saccharomyces cerevisiae. Sci Rep 8:11919. 10.1038/s41598-018-30305-z

59. Alexandrov LB, Kim J, Haradhvala NJ, et al (2020) The repertoire of mutational signatures in human cancer. Nature 578:94–101. 10.1038/s41586-020-1943-3

60. Bohnert KA, Gould KL (2012) Cytokinesis-based constraints on polarized cell growth in fission yeast. PLoS Genet 8:e1003004. 10.1371/journal.pgen.1003004

61. Kippert F, Lloyd D (1995) The aniline blue fluorochrome specifically stains the septum of both live and fixed Schizosaccharomyces pombe cells. FEMS Microbiol Lett 132:215–219. 10.1111/j.1574-6968.1995.tb07836.x

62. Duong H-Q, Hong YB, Kim JS, et al (2013) Inhibition of checkpoint kinase 2 (CHK2) enhances sensitivity of pancreatic adenocarcinoma cells to gemcitabine. J Cell Mol Med 17:1261–1270. 10.1111/jcmm.12101

63. Höfer S, Frasch L, Brajkovic S, et al (2025) Gemcitabine and ATR inhibitors synergize to kill PDAC cells by blocking DNA damage response. Mol Syst Biol 21:231–253. 10.1038/s44320-025-00085-6

64. Kianfard Z, Cheung KJ, Patel PN, et al (2025) Genetic profiles of gemcitabine sensitivity in Schizosaccharomyces pombe. in prep:

65. Dixon SJ, Andrews BJ, Boone C (2009) Exploring the conservation of synthetic lethal genetic interaction networks. Commun Integr Biol 2:78–81

66. Brown JS, O’Carrigan B, Jackson SP, Yap TA (2017) Targeting DNA Repair in Cancer: Beyond PARP Inhibitors. Cancer Discov 7:20–37. 10.1158/2159-8290.CD-16-0860

67. Fordham SE, Blair HJ, Elstob CJ, et al (2018) Inhibition of ATR acutely sensitizes acute myeloid leukemia cells to nucleoside analogs that target ribonucleotide reductase. Blood Adv 2:1157–1169. 10.1182/bloodadvances.2017015214

68. Bradbury A, Hall S, Curtin N, Drew Y (2020) Targeting ATR as Cancer Therapy: A new era for synthetic lethality and synergistic combinations? Pharmacol Ther 207:107450. 10.1016/j.pharmthera.2019.107450

69. Lynx MD, Bentley AT, McKee EE (2006) 3’-Azido-3’-deoxythymidine (AZT) inhibits thymidine phosphorylation in isolated rat liver mitochondria: a possible mechanism of AZT hepatotoxicity. Biochem Pharmacol 71:1342–1348. 10.1016/j.bcp.2006.01.003

70. Blanco F, García-Benayas T, José de la Cruz J, et al (2003) First-line therapy and mitochondrial damage: different nucleosides, different findings. HIV Clin Trials 4:11–19. 10.1310/HF1J-3P6K-1K9H-AGPY

71. Sutera VA Jr, Lovett ST (2006) The role of replication initiation control in promoting survival of replication fork damage. Mol Microbiol 60:229–39

72. Canadian Cancer Statistics Advisory in collaboration with, the Canadian Cancer Society, Statistics Canada and the, Public Health Agency of Canada (2022) Canadian Cancer Statistics: A 2022 special report on cancer prevalence

73. Bukhari AB, Chan GK, Gamper AM (2022) Targeting the DNA Damage Response for Cancer Therapy by Inhibiting the Kinase Wee1. Front Oncol 12:828684. 10.3389/fonc.2022.828684

74. Fu S, Yao S, Yuan Y, et al (2023) Multicenter Phase II Trial of the WEE1 Inhibitor Adavosertib in Refractory Solid Tumors Harboring CCNE1 Amplification. J Clin Oncol Off J Am Soc Clin Oncol 41:1725–1734. 10.1200/JCO.22.00830

75. Cuneo KC, Morgan MA, Sahai V, et al (2019) Dose Escalation Trial of the Wee1 Inhibitor Adavosertib (AZD1775) in Combination With Gemcitabine and Radiation for Patients With Locally Advanced Pancreatic Cancer. J Clin Oncol Off J Am Soc Clin Oncol 37:2643–2650. 10.1200/JCO.19.00730

76. Mueller S, Cooney T, Yang X, et al (2022) Wee1 kinase inhibitor adavosertib with radiation in newly diagnosed diffuse intrinsic pontine glioma: A Children’s Oncology Group phase I consortium study. Neuro-Oncol Adv 4:vdac073. 10.1093/noajnl/vdac073

77. Zhang C, Peng K, Liu Q, et al (2024) Adavosertib and beyond: Biomarkers, drug combination and toxicity of WEE1 inhibitors. Crit Rev Oncol Hematol 193:104233. 10.1016/j.critrevonc.2023.104233

78. Sabatinos SA, Forsburg SL (2010) Molecular genetics of Schizosaccharomyces pombe. Methods Enzymol 470:759–95. 10.1016/S0076-6879(10)70032-X

79. Petersen J, Russell P (2016) Growth and the Environment of Schizosaccharomyces pombe. Cold Spring Harb Protoc 2016:pdb.top079764. 10.1101/pdb.top079764

80. Karam E, Sabatinos SA (2025) Investigating Fission Yeast Mutagenesis Using Canavanine Sensitivity Assays. Methods Mol Biol Clifton NJ 2862:195–208. 10.1007/978-1-0716-4168-2_14

81. Schindelin J, Arganda-Carreras I, Frise E, et al (2012) Fiji: an open-source platform for biological-image analysis. Nat Methods 9:676–682. 10.1038/nmeth.2019

