## Supplementary Figure S1 for "Checkpoint-dependent sensitivities to nucleoside analogues uncover specific patterns of genomic instability"

SUPPLEMENTARTY DATA FILE
for

### Checkpoint-dependent sensitivities to nucleoside analogues uncover specific patterns of genomic instability

Zainab Kagalwala[1,2,‡], Mohammed Ayan Chhipa[1,3,‡], Zohreh Kianfard[1,3], Essam Karam [1,3], Sirasie P Magalage[1,3], and Sarah A Sabatinos[1,3,*]

**Contents:**

S1: Supplementary Figure S1. Insensitivity to FUdR increases above the IC50 dose of BrdU.

Supplementary Figure S1:

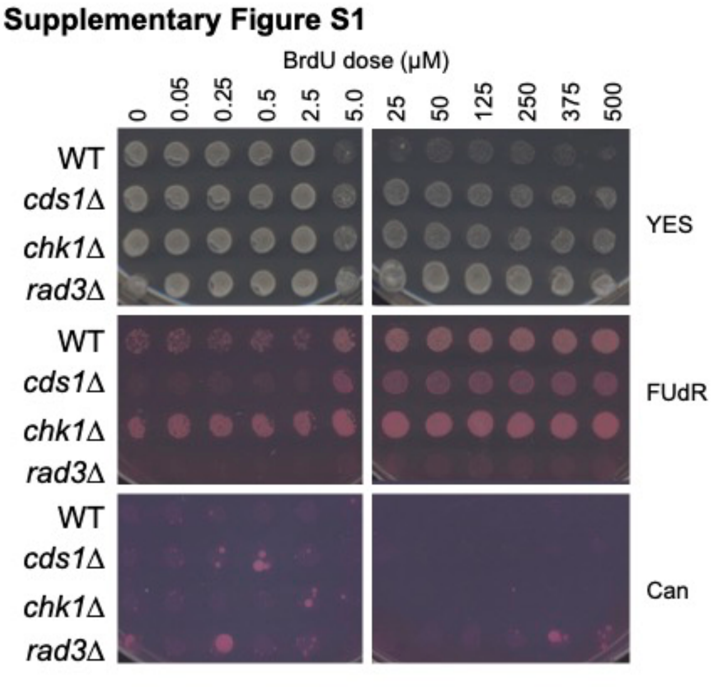


**Supplementary Figure S1. Insensitivity to FUdR increases above the IC50 dose of BrdU.** Shown on top are the YES plates after 48 h exposure to varying doses of BrdU in PMG medium with supplements. Concurrently, treated cells were also pinned onto FUdR (middle) to assess HSV-TK function. Cells were also pinned onto PMG-HULA medium containing 70 μg/mL canavanine sulfate [80]. Canavanine (Can) can be used to detect forward mutation in the canavanine sensitivity pathway. Can mutants are large colonies and are more common in *cds1*Δ, *chk1*Δ and *rad3*Δ up to the IC50 dose. Phloxine B was added to FUdR and Can plates to assess cell health; darker pink staining means that cells are less able to excrete the dye and the spots are darker pink.
